# Aromatic and arginine content drives multiphasic condensation of protein–RNA mixtures

**DOI:** 10.1101/2023.04.05.535720

**Authors:** Pin Yu Chew, Jerelle A. Joseph, Rosana Collepardo-Guevara, Aleks Reinhardt

## Abstract

Multiphasic architectures are found ubiquitously in biomolecular condensates and are thought to have important implications for the organisation of multiple chemical reactions within the same compartment. Many of these multiphasic condensates contain RNA in addition to proteins. Here, we investigate the importance of different interactions in multiphasic condensates comprising two different proteins and RNA using computer simulations with a residue-resolution coarse-grained model of proteins and RNA. We find that in multilayered condensates containing RNA in both phases, protein–RNA interactions dominate, with aromatic residues and arginine forming the key stabilising interactions. The total aromatic and arginine content of the two proteins must be appreciably different for distinct phases to form, and we show that this difference increases as the system is driven towards greater multiphasicity. Using the trends observed in the different interaction energies of this system, we demonstrate that we can also construct multilayered condensates with RNA preferentially concentrated in one phase. The ‘rules’ identified can thus enable the design of synthetic multiphasic condensates to facilitate further study of their organisation and function.

## I. INTRODUCTION

The formation of biomolecular condensates by phase separation achieves spatio-temporal organisation within the cell by creating specific intracellular environments that help to regulate cellular activity such as cell signalling,^1–4^ stress response,^5–9^ regulation of transcription,^10–15^ and associated chemical reactions.^16,17^ These condensates often comprise numerous components, and sometimes compositional heterogeneity can be seen within the same compartment. Such ‘multiphasic’ core–shell architectures have, for example, been observed in condensates inside cells, such as the nucleolus,^18,19^ P granules^20–22^ and stress granules.^5^ The internal subcompartmentalisation of biomolecules to form multiphasic structures may be a possible way by which cells can segregate different biological processes.^18,19,23–26^ Simple model systems with a minimal number of components have been used to study the properties of condensates with multiphasic architectures.^27–31^ Moreover, relatively simple multiphasic systems have been designed to segregate enzymes and form an enzymatic cascade network of reactions,^32^ and to direct the flow of biochemical reactions such as by coupling in-vitro transcription and translation across different subcompartments within the same condensate.^33^

Computer simulations are a useful tool to help elucidate the underlying molecular interactions governing the complex phase behaviour of biomolecular mixtures. All-atom,^34–41^ residue-resolution,^42–49^ and minimal ‘stickers-and-spacers’ and patchy-particle^50–56^ protein models can all be used to gain ‘ different insights into the driving forces of condensate assembly.

Simulations of biomolecular phase separation have also been combined with theory^57–61^ and with other computational approaches, such as machine-learning methods to predict properties and to parameterise force fields,^46–48,62–64^ and evolutionary algorithms to promote or inhibit phase separation or a desired i spatial organisation of biomolecules inside condensates.^65,66^

Despite the prevalence of biomolecular multiphasic condensates, the molecular drivers governing their thermodynamic i immiscibility are not very well understood. Weak multivalent attractive interactions are usually required for phase separation.^4,51,54,58,67–71^ Multiphasicity has been shown to emerge when there are competing interactions for a shared binding partner and the overall interaction strengths in one phase are sufficiently different from that of another phase for them to be immiscible.^18,27–31,66,72,73^ One intriguing observation is that in i all the multiphasic condensates observed in cells thus far, not only proteins are present, but also RNA.^74–76^ In protein–RNA condensates, the relevant stabilising interactions that RNA nucleobases can form include associative electrostatic interactions with positively charged residues such as lysine (K) and arginine (R), π–π stacking interactions with aromatic residues such as tyrosine (Y) and phenylalanine (F), as well as cation-π interactions with arginine. These interactions all contribute to different extents to the stabilisation of the condensate, and the type of interaction that dominates for a given protein/RNA sequence may be responsible for inducing the compositional demixing that results in multiphasic architectures. More specifically, it has been shown that arginine is especially abundant in RNA-binding proteins.^75,76^ As arginine is positively charged, it can interact with RNA with a direct Coulomb interaction as well as a cation–π interaction. Interestingly, even though arginine and lysine both comprise an alkyl side chain with nitrogen functionalisation, and are positively charged at physiological pH, they are well known to be unequal contributors to biomolecular phase separation.^27,39,52,54,77^ The cation–π interactions established by arginine with aromatic rings (such as RNA bases) are not only stronger than those formed by lysine,^78^ but also less sensitive to screening by counterions in solution.^39,52,79^ Such unequal behaviour of arginine versus lysine has been explained by the higher hydrophobicit^27,39,80,81^ and the less favourable free energy of hydration of arginine,^82^ and the hybrid cation–π/π–π nature of the bonds arginine establishes with π-rich species.^39,83^ Indeed, in several studies, mutating arginine to lysine in protein sequences greatly destabilised condensates,^77,84^ whilst increasing the proportion of arginine favoured phase separation^.44,85^ Whether arginine plays a similarly important role also in multiphasic condensates of mixtures of proteins and RNA is an open question that we explore in this manuscript. The crucial role of RNA in stabilising biomolecular condensates has been identified as one of the glaring omissions in the field,^86^ and we aim to begin to address it here.

## II. RESULTS

In order to understand what kinds of interaction are likely to play an important role in maintaining the multiphasicity of protein/RNA condensates, we first investigate the behaviour of simple model systems that contain RNA and that have been observed experimentally to form multiphasic condensates. One example of such a mixture contains equimolar amounts of poly-arginine and poly-lysine, together with a charge-matched amount of poly-uracil to bring the net charge of the system to zero. Such a charge-neutral system of poly-arginine, polylysine and poly-uracil has been observed to form multilayered droplets in *in-vitro* experiments,^27^ with the poly-arginine-rich phase being concentrated in the core of the condensate whilst the outer layer is enriched in poly-lysine. Systems of both poly-arginine and poly-lysine can also separately form condensates with RNA, but with different interfacial free-energy densities and critical temperatures.^27^ We can reproduce such multiphase compartmentalisation [Fig. 1] in our simulations using Mpipi, a residue-solution coarse-grained model that can predict the critical solution temperature of protein solutions in good agreement with experiment.^52^

**Figure 1.**
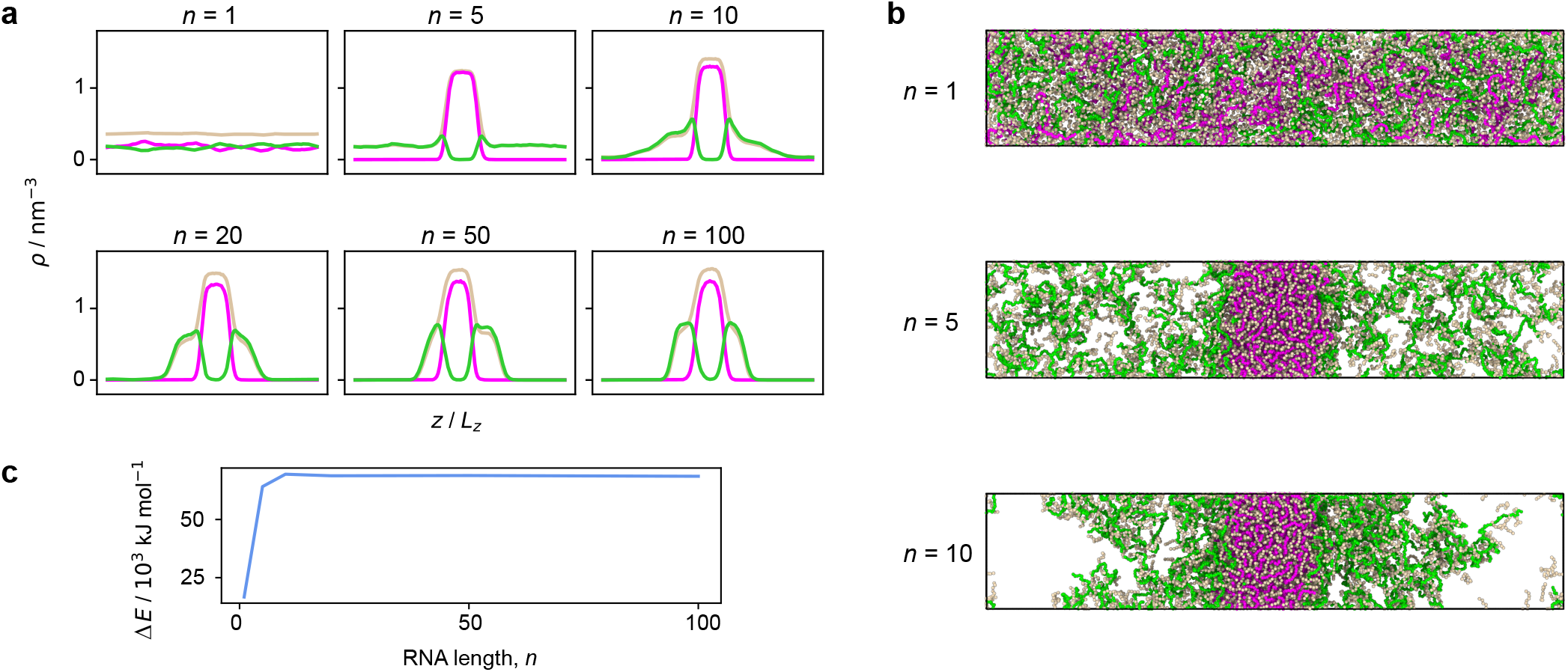
**a** Variation in the density profiles of a mixture of an equimolar amount of poly-arginine, R_50_, and poly-lysine, K_50_, with a charged-matched amount of poly-uracil, U*_n_*, with the length of the RNA polymer, *n*. **b** Simulation snapshots of the systems in **a** with *n* = 1, 5 and 10. Multilayered droplets with the arginine-rich phase at the centre and the lysine-rich phase on the outside form with poly-uracil of length *n* > 10. **c** Difference in the interaction energies of arginine–uracil (R–U) and lysine–uracil (K–U), as a function of *n*. Δ*E* is the difference in the total R–U and K–U interaction energies summed over all the beads in the system, averaged over time.

Since solutions of poly-arginine and poly-lysine on their own, as well as mixtures of poly-arginine and poly-lysine, do not phase separate in the absence of RNA in solution conditions approaching those inside cells, e.g. 0.15 m NaCl in the absence of crowders, we might expect an important contribution to the stabilising interactions of this system that enable phase separation to arise from the electrostatic attraction between the positively charged arginine/lysine residues and the negatively charged RNA nucleotides. Indeed, the inability of poly-arginine and/or poly-lysine to condense without RNA does in part result from the electrostatic repulsion between the positively charged residues. However, if we set the charge on arginine and lysine to be zero in our simulations, crudely mimicking relatively high salt concentrations where the screening length is small,^87^ zero-charge-poly-arginine and/or zero-charge-poly-lysine are still unable to form condensates without RNA. The inability of poly-lysine and poly-arginine to phase separate without RNA thus appears to arise from a combination of the destabilisation from the electrostatic repulsion amongst the positive charges and a lack of other sufficiently stabilising attractive interactions.

We also observe in our simulations that the phase behaviour is dependent on the length of the RNA polymers [Fig. 1], which is consistent with observations from experiments and simulations of similar systems which phase separate via complex coacervation.^27,88^ An equimolar system of poly-arginine R_50_ and poly-lysine K_50_ that is made charge-neutral by the addition of RNA mononucleotides also does not form stable condensates; charge neutralisation is therefore not sufficient for phase separation, and we require the RNA to have a minimum length, and hence a sufficient valency, to ensure the formation of the percolated network of inter-molecular bonds that sustains the condensate.^51,86,89,90^ If RNA nucleotides form longer polymer chains, the resulting entropic penalty of forming a condensed phase is likely to be lower; this is consistent with the Flory–Huggins theory of polymer solutions, where the critical point increases with polymer length.^91^ Moreover, a longer RNA strand can increase the density of connections in the condensed phase, since the negatively charged RNA bases are forced to remain close to one another by being covalently bonded.^88^ Lysine–uracil interactions are weaker than arginine–uracil interaction, resulting in a weaker enthalpic favourability for demixing. For phase separation to ensue, this weaker enthalpic driving force must therefore be counteracted by a better connected liquid network, and the minimum required length of RNA to form the poly-lysine K_50_ phase is thus longer. In the rest of this work, we use poly-uracil U_10_ as the appropriate RNA counterpart to the proteins we consider, since it is the shortest length that enables both R_50_ and K_50_ to phase separate, and the difference between between the arginine–RNA and lysine–RNA interaction energies plateaus at a length of approximately 10 nucleotides per RNA molecule [Fig. 1**c**]. It is convenient to investigate the behaviour of systems with RNA chains that are as short as possible while resulting in multiphasic behaviour, since longer RNA chains require considerably longer equilibration times in computer simulations.

To investigate which interactions are responsible for driving the formation of multilayered protein–RNA condensates, we can start from a phase-separated system of high multiphasicity and gradually evolve it towards decreasing multiphasicity where the two phases become more similar in composition^66^ using a genetic algorithm (see Methods). By exploring the difference in composition of sequences exhibiting low and high multiphasicity, we can obtain physical insight into the principal driving forces behind multiphasic behaviour, and in turn gain an intuition for how best to design initial sequences of mixtures for possible target applications. Such an approach has already proved fruitful in previous work on single-component^65^ and multi-component^66^ protein systems. To be able to use a genetic algorithm, we must first define a function that allows us to quantify the degree of ‘multiphasicity’ of a condensate. Although multiphasic condensates can exist in various architectures, here we are interested in multiphasic systems that correspond to a phase-separated system where the condensed phase forms two distinct layered phases,^66^ each with a different composition. In an elongated simulation box, the disfavourable interfacial free energy is minimised by planar interfaces.^66^ If a multiphasic system forms, we thus expect to have a layered condensate, with a vapour-like phase on the outside, followed by the condensed phase with the lower interfacial tension with the vapour,^92,93^ and the second condensed phase at the centre [Fig. 1**b**]. In order to favour the evolution towards less multiphasic condensed phases, we use the fitness function

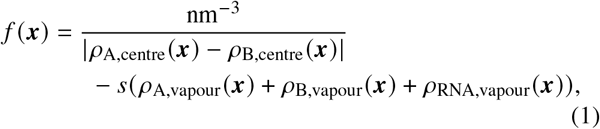

where *ρ_A_*(***x***) and *ρ_B_*(***x***) are averaged number densities of the two different protein sequences A and B in the mixture. *ρ_i,centre_*(***x***) and *ρ_i,vapour_*(***x***) denote the number density of species *i* in the core of the multilayered condensate and in the dilute phase, respectively. The first term in the fitness function is the reciprocal of the difference in densities of the two protein species; the larger this term, the more homogeneous the two coexisting phases become in terms of the distribution of the two proteins. To try to ensure that a stable condensate still forms, the second term penalises the accumulation of any species in the dilute phase. We scale this penalty term by a weighting parameter *s*, and unless otherwise stated, we use *s* = 5 nm^3^ in all genetic-algorithm runs to provide a reasonable balance between penalising the accumulation of molecules in the dilute phase and the ability of the molecules to mix in the condensed phase. By contrast, to drive the overall mixture towards increasing multiphasicity, we can use the fitness function

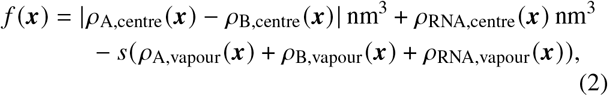

where *s* again penalises the accumulation of any species in the vapour phase. Similar fitness functions involving the densities of the different species in the vapour phase and inner layer have been used to study multilayered droplets of proteins in the absence of RNA in previous work.^66^ Other order parameters have also been used to quantify such multiphasic compartmentalisation, e.g. looking at the compositional asymmetry of the two dense phases^59^ or the intra- and inter-species pair correlation functions as a function of intermolecular separation.^60,61^ These fitness functions have been shown to correlate well with the simple density-based fitness function proposed above.^66^

Having introduced a suitable fitness function, we start genetic-algorithm runs from an initial system comprising an equimolar amount of R_50_ and K_50_ with a corresponding charge-matched amount of U_10_. This system spontaneously forms an initial multilayered condensate with high multiphasicity in our simulations and in in vitro experiments.^27^ Using the fitness function of Eq. (1), we consider three scenarios: in independent simulations, we (a) evolve the sequence of K_50_ only whilst keeping R_50_ unchanged throughout, (b) evolve the sequence of R_50_ only whilst keeping K_50_ unchanged throughout, and (c) evolve both K_50_ and R_50_ simultaneously with the same mutation rate applied to both sequences [Fig. 2**a–c**]. In cases (a) and (c), we are able to obtain a stable homogeneous condensate, whereas in case (b), where we evolve only R_50_, a stable homogeneous condensate does not form, even if we increase the weighting of the penalty term disfavouring full mixing. Specifically, increasing the value of the parameter *s* in our fitness function yields either a well-mixed, non-phase-separated fluid (with *s* = 10^6^ nm^3^, Fig. 2**b**), or the system remains in a multiphasic state with the evolved sequence not changing significantly from its initial sequence (with *s* = 10^8^ nm^3^, Fig. S6). Since in this case the starting protein sequence contains only arginines, all mutations inevitably involve replacing arginine with other residues throughout the evolution. These results thus suggest that arginine stabilises protein–RNA condensates: a certain fraction of arginine in the condensate seems to be necessary to maintain its stability in the presence of RNA. Interestingly, although the arginine fraction of what was initially R_50_ decreases both when we evolve solely R_50_ [Fig. 2**b**] and when we evolve both proteins [Fig. 2**c**], the final state is markedly different: whilst the degree of multiphasicity decreases in both cases, in the case where both sequences are evolved, the decrease of arginine content in one sequence is compensated by an increase in arginine and aromatic residues in the other sequence, which seems to be necessary to enable a monophasic condensate to form. Both arginine and aromatic residues thus appear to be important when RNA is present, since they can form strong attractive interactions with the nucleobases.

**Figure 2.**
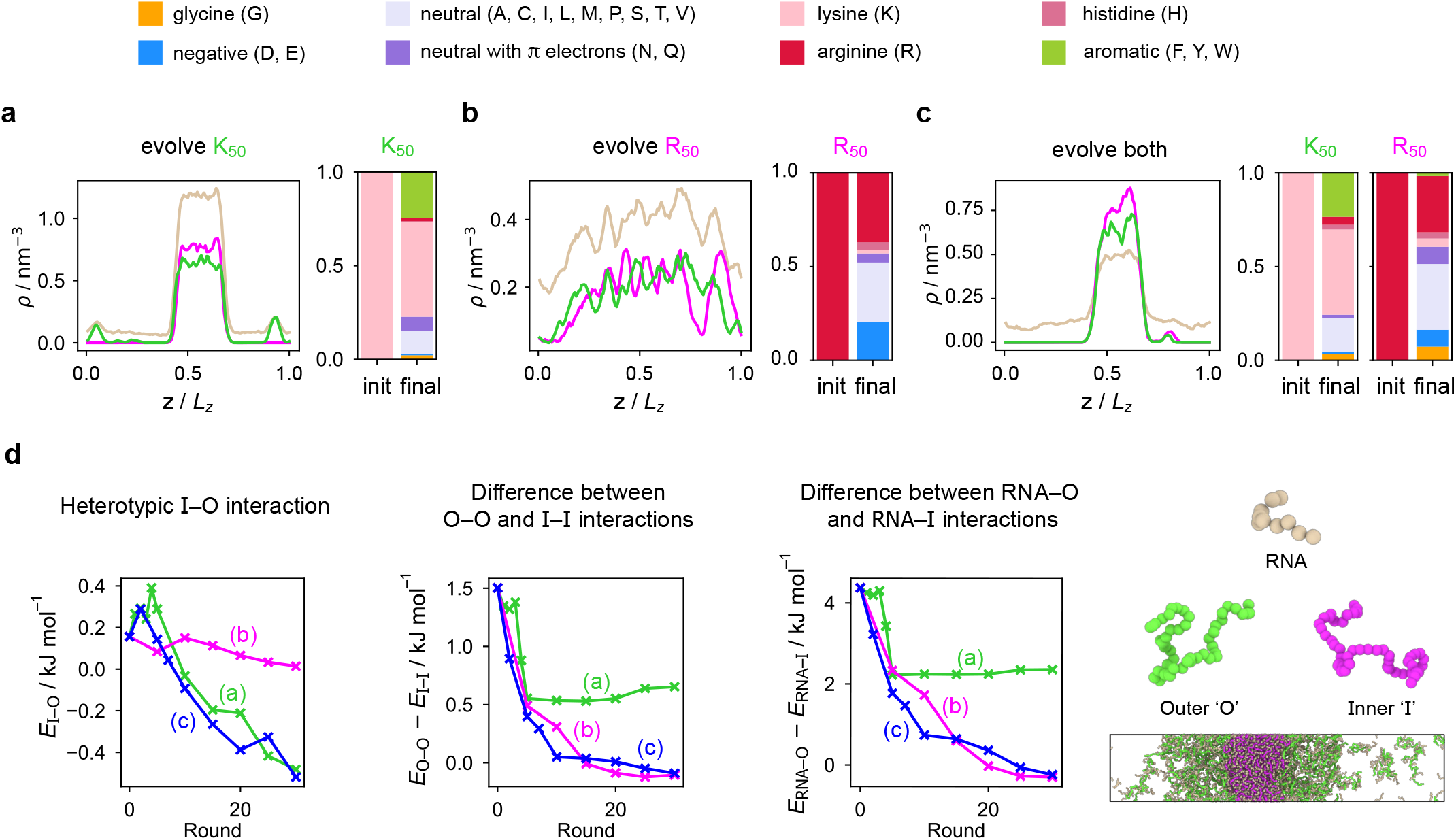
Density profiles of the final evolved system with maximum fitness and changes in composition of the evolved sequence in the genetic-algorithm runs towards decreasing multiphasicity, where we evolve **a** K_50_ or **b** R_50_ in separate runs, or **c** both sequences simultaneously from the initial starting system in Fig. 1**a** with *n* = 10. The pink, green and light brown curves correspond to the density profiles of (evolved) R_50_, (evolved) K_50_ and U_10_ respectively. In all three cases, the final composition, averaged across all 20 sequences in the population of the final round, is shown in terms of amino-acid types, according to the key at the top of the figure. **d** Interaction energies between the different species within the condensate as a function of the round number in the genetic-algorithm run. Green, magenta and blue curves correspond to the cases in **a**, **b** and **c** respectively, as labelled. ‘O’ and ‘I’ refer to the proteins concentrated in the outer and inner phases of the multilayered condensates formed.

As we have already discussed, single-component systems of either R_50_ or K_50_ are not able to form stable condensates on their own at salt concentrations close to physiological conditions, and RNA is required for charge neutralisation. Although such charge neutralisation is not sufficient to drive demixing, and the added RNA must also be of a certain minimum length, the electrostatic interactions between the positively charged arginine/lysine and the negatively charged uracil are clearly important for controlling phase behaviour. However, despite both arginine and lysine being positively charged at physiological pH, there are notable differences in the roles of the two amino acids. In our simulations, phase-separated condensates of the protein R_*m*_K_50-*m*_, where *m* ∈ ℤ is varied between 0 and 50, mixed with a charge-matched amount of U_10_, are progressively more dense [Fig. S7] and hence more stable as the protein contains more arginine than lysine (i.e. as *m* increases), again suggesting that arginine plays an outsize role in phase stability. Both arginine and lysine can form electrostatic interactions and cation–π interactions with the RNA nucleotides; however, only arginine can form π–π interactions between the guanidinium group of the arginine side chain and the RNA nucleotide.^36,85,94^ These differences between arginine and lysine are described implicitly in the Mpipi residue-resolution coarse-grained model that we use,^52^ which assigns stronger short-range interactions to the arginine–uracil pair than to the lysine–uracil pair, enabling simulations to capture the difference in thermodynamic behaviour between the two residues.

More strikingly, our simulations also reveal an important role of arginine in the modulation of RNA–protein multiphasic condensates that display an inner phase surrounded by an outer phase of different compositions. Specifically, we consider three-component RNA–protein multiphase condensates made of two different types of proteins (namely R_*m*_K_50-*m*_ and R_50-*m*_K_*m*_, where 0 ≤ *m* ≤ 50) and a charge-matched amount of RNA. These mixtures form multiphasic condensates with an ‘inner’ and an ‘outer’ phase. We refer to the majority-component proteins in these two regions as protein I (‘inner’) and protein O (‘outer’), respectively. We find that the protein that has a higher arginine content is always in the inner phase, i.e. protein I is R_*m*_K_50-*m*_ if *m* ≥ 25 and R_50-*m*_K_*m*_ otherwise. Moreover, the greater the difference in arginine composition between the two sequences, the greater the multiphasicity of the resulting condensate when quantified with the fitness function of Eq. (2) [Fig. 3].

**Figure 3.**
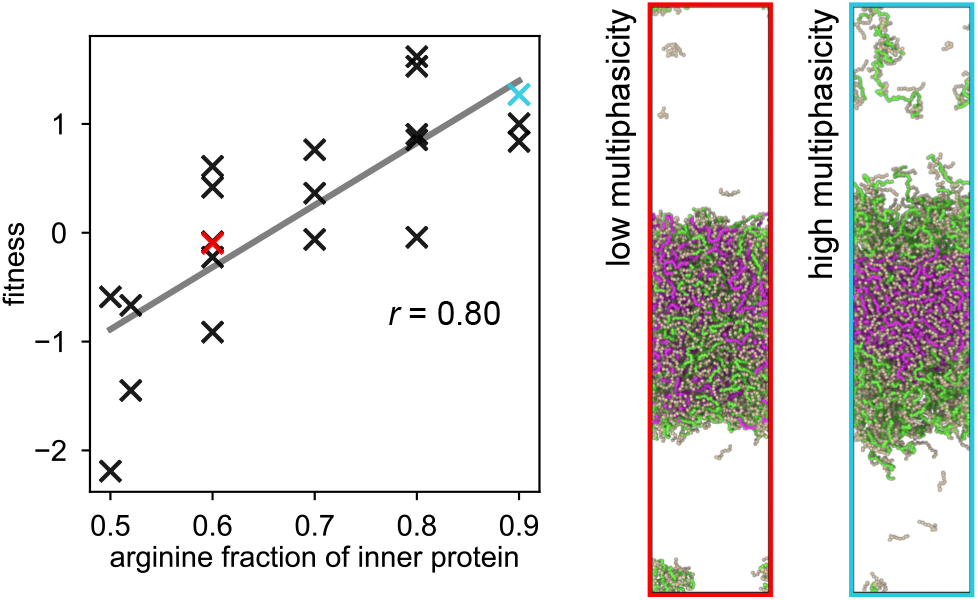
Variation in the degree of multiphasicity, as quantified by the fitness function in Eq. (2), with the fraction of arginine in the inner protein sequence in three-component RNA–protein multiphase condensates made up of equimolar amounts of R_*m*_K_50-*m*_ and R_50-*m*_K_*m*_ (where 0 ≤ ***m*** ≤ 50), and a charge-matched amount of RNA. The total arginine and lysine content is kept fixed across both protein species. The multiple points with the same arginine composition correspond to sequences with different patterning of the arginine and lysine residues. The two simulation snapshots provided as examples of low and high multiphasicity correspond to the points on the graph with the same colour as the box outline. There are small variations in multiphasicity depending on the patterning, but the degree of multiphasicity broadly increases with the increasing fraction of arginine in the inner sequence.

Although the three genetic-algorithm runs towards reduced multiphasicity [Fig. 2**a–c**] are more complex because residues other than arginine and lysine are present, the same observation holds: the degree of multiphasicity is reduced when the difference in the proportion of residues that form the stabilising attractive interactions with RNA (i.e. arginine and aromatic residues) between the two proteins is lower. When only K_50_ is evolved, the residues in K_50_ are replaced with more strongly interacting residues, as approximated by the *ε_i_* value in the Mpipi model. The replaced residues mostly end up as aromatic residues (Y, F and W), and make up about a quarter of the final evolved sequence at the end of the run [Fig. 2**a**, Fig. S5**a**], whilst the partner sequence is still R_50_. By contrast, when only R_50_ is evolved, the residues in R_50_ are replaced with more weakly interacting residues [Fig. 2**b**, Fig. S5**b**]. In the run where we evolve both sequences simultaneously, both sequences evolve in similar ways to how the individual sequences evolved separately [Fig. 2**c**, Fig. S5**c**]. In all cases, the degree of multiphasicity is thus reduced as the two proteins become more similar in terms of overall residue interaction strengths.

In all three cases, the average net charge of the evolved sequences decreases as the positively charged lysine and arginine residues are replaced with neutral or negatively charged ones [Fig. S5]. The greatest decrease in average net charge of the evolved sequence occurs for the case where only R_50_ is evolved [Fig. 2**b**, 〈Δ〉_(b)_ = −0.56*e* compared to 〈Δ〉_(a)_ = −0.32*e* and 〈Δ〉_(c)_ = −0.42*e*], with a large proportion of residues being exchanged with negatively charged residues (D and E) over neutral ones. Since the initial mixture is overall charge neutral, this large decrease in the net charge of the evolved protein results in the mixture becoming negatively charged overall. An excess negative charge contributes to the destabilisation of the RNA– protein condensate;^95,96^ however, our simulations indicate that other factors are also at play: when we set the charge of a fraction of the negatively charged species to zero to recover a mixture with zero overall net charge, the system is still unable to undergo phase separation. Hence we can conclude that attractive electrostatic and short-ranged interactions are evidently not sufficient in the first place to effect phase separation in this case. In other words, replacing arginine with other residues that interact less strongly overall results in the inability to undergo phase separation, which further underlies the importance of arginine in protein–RNA condensates.

Finally, to investigate how these changes in protein composition affect the relative contributions of the different interactions between the various components in the mixture, we analyse the changes in the different contributions from the homo- vs heterotypic interaction energies between the different components as the systems are evolved towards lower multiphasicity [Fig. 2**d**]. As before, we label the two proteins as protein I, which was originally at the centre of the multiphasic condensate (and started off as R_50_), and protein O, which was originally in the outer condensed phase (and started off as K_50_). In all cases, the homotypic interactions, I–I and O–O, become less repulsive on average, indicating that the positively charged residues are being exchanged for neutral or negatively charged ones, with the magnitude of the difference between the I–I and O–O energies decreasing. The heterotypic I–O interactions also become less repulsive in both cases, and the I–O interaction can eventually even become attractive. Since RNA acts as a glue holding the multiphasic condensate together, the heterotypic I–RNA and O–RNA interactions are the attractive interactions driving phase separation in each individual liquid-like phase, as discussed previously and seen in their relative magnitudes compared to the protein–protein interactions. It is the change in these interactions that appears to have the greatest effect on the overall phase behaviour. In all cases, the I–RNA interactions become less attractive as the systems become less multiphasic, while the O–RNA interactions become more attractive, with the mean difference between I–RNA and O–RNA decreasing in magnitude [Fig. 2**d**]. It is well-known that compositional demixing can occur when the difference in interaction strengths between the components within each of the immiscible phases is sufficiently large to maintain multiphasicity;^18,27,29–31,66,73^ for the case of protein–RNA condensates, our observations suggest that it is the difference between the two different protein–RNA interaction energies that must be large for such multiphasic condensates to be stabilised.

In order to check the robustness of these observations, we can determine not only what happens as the systems are evolved towards a monophasic condensed phase, but also if the reverse process is feasible. To this end, we begin with a multi-component monophasic condensate [Fig. 4**a**] obtained when evolving the R_50_/K_50_/U_10_ system towards lower multiphasicity by evolving both the inner and the outer protein sequence, i.e. corresponding to case (c) above, where the inner protein was evolved from R_50_ and the outer protein from K_50_. We then evolve it towards higher multiphasicity by evolving both protein sequences in the system simultaneously, using the fitness function of Eq. (2). The final system in Fig. 4**a** displays two phases in a multilayered arrangement, with one protein preferentially concentrated in either phase. The changes in composition are the opposite to what we observe in Fig. 2**c**: when the system is evolved towards increasing multiphasicity, the difference in the fraction of residues that form the stabilising attractive interactions with RNA (i.e. arginine and aromatic residues) between the two proteins increases, with the inner protein having a larger proportion of aromatic residues and arginine [Fig. 4**b,c**]. To confirm the role of arginine and aromatic residues in driving compositional demixing in these protein–RNA condensates, we perform three additional genetic-algorithm runs where (a) mutations to arginine are not allowed, (b) mutations to aromatic residues are not allowed, and (c) mutations to both arginine and aromatic residues are not allowed. The resulting changes in composition [Fig. **S8a–c**] confirm that the difference in the combined fraction of arginine and aromatic residues between the inner and outer protein always increases, and this increase can be achieved by increasing either type of residue in the inner protein or decreasing either in the outer protein. In case (c) where mutations to both arginine and aromatic residues are not allowed, the evolution towards higher multiphasicity is much less successful. Finally, we note that when the system is evolved towards increasing multiphasicity, the corresponding changes in interaction energies [Fig. S8**e**] are the opposite of those shown in Fig. 2**d** for evolution towards decreasing multiphasicity.

**Figure 4.**
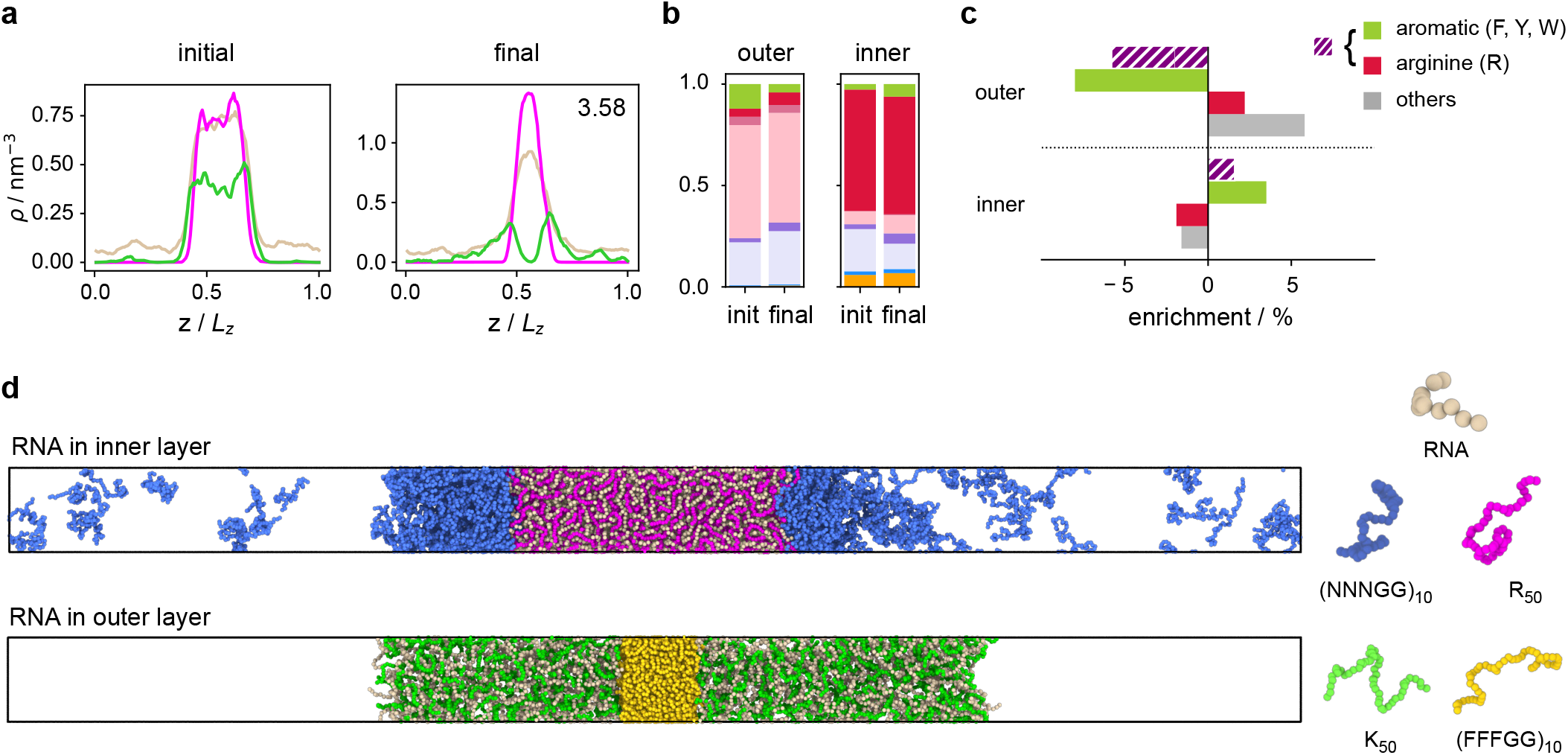
Evolution towards higher multiphasicity. **a** Density profiles of the initial and final systems in the genetic-algorithm run where we evolve both protein sequences simultaneously towards increasing multiphasicity of the system. In the top right-hand corner of the density plot of the final system, we give the fitness value of the final system relative to the initial system. **b** Changes in composition of the outer and inner sequences. For both sequences, the final composition is averaged across the all 20 sequences in the population of the final round. In particular, we highlight the enrichment of arginine and/or aromatic residues in **c**. The enrichment in this case is defined as the difference in fraction of the residue type of interest between the final and initial sequence. We observe that the difference in the fraction of arginine and aromatic residues increases between the outer and inner sequence as we evolve towards increasing multiphasicity. **d** Multilayered systems with simple generic protein sequences with RNA preferentially concentrated in one phase. RNA in inner layer: R_50_+U50 in the centre and (NNNGG)10 on the outside. RNA in outer layer: (FFFGG)_10_ in the centre and K_50_+U50 on the outside.

In the systems we have discussed so far, the RNA molecules (U_10_) are distributed evenly across both dense phases in the multiphasic condensate. However, in such condensates inside cells, RNA is sometimes concentrated preferentially in different phases, which enables different processing of the RNA molecules depending on their function or cellular environment, and it would be useful to understand how cells can make use of the different interactions occurring in the different phases to segregate RNA. As a proof of concept, we thus design multilayered condensates with RNA concentrated preferentially in one phase and depleted in the other. It may initially seem that a straightforward way to achieve this would be to design a suitable fitness function that favours the unequal partitioning of RNA across the two condensed phases and evolve one of the multiphasic systems we have already investigated. However, in reality, this would be a rather difficult process, since one of the two coexisting proteins in the final multiphasic architecture would need to form a condensed phase on its own, without the involvement of RNA. As we have already discussed, neither of the two protein species we have considered so far is able to undergo demixing. To get a condensed phase without RNA using a genetic-algorithm approach, one of the proteins would thus have to evolve to one that is able to undergo phase separation by itself. Although this is not impossible, starting from a protein that is very far in sequence space from the ‘target’ one would entail large changes to the sequence during the course of the evolution, which would render the process inefficient. Instead, we can use the lessons learnt about the dominant interactions that drive phase separation to design simple systems with RNA preferentially concentrated in one phase by hand. The total interaction strength within the outer phase should be significantly weaker than within the inner phase for a multilayered condensate to form.^66^ In the Mpipi model, aromatic–aromatic interactions are more favourable than aromatic–RNA interactions, and these in turn are more favourable than arginine–RNA interactions. If we wish for RNA to be concentrated in the inner layer and depleted from the outer layer, the protein sequence on the outside should be able to phase separate by itself, but ought not to contain a significant amount of aromatic residues, since such residues favour mixing with the RNA on the inside, given that aromatic–RNA interactions are stronger than arginine–RNA ones. By contrast, if RNA is to be on the outside and depleted from the inner layer, the inner protein sequence should be rich in aromatic residues, since aromatic–aromatic interactions are stronger than aromatic–RNA, and so it would be energetically more favourable for such a sequence to form a phase by itself and exclude RNA. We show examples of such systems in Fig. 4**d**. From these initial systems, we could then use a coevolution approach^66^ with a modified fitness function to drive the RNA to the specific region into which we wish to partition it.

## III. DISCUSSION

In this work, we use our residue/nucleotide-resolution coarse-grained model for proteins and disordered RNA to investigate the role of amino-acid sequence composition in driving the multiphasicity of protein–RNA condensates with direct-coexistence molecular-dynamics simulations and a genetic algorithm. Our simulations indicate that the formation of multi-component multilayered condensates made up of two distinct proteins and RNA is principally determined by the proteins’ aromatic and arginine composition. Specifically, the degree of multiphasicity increases with the difference in fraction of arginine and aromatic residues between the two proteins in the condensate. The strong impact of arginine and aromatics in the multiphasicity of protein-RNA condensates is tightly linked to (a) the unrivalled ability of such residues to establish strong attractive interactions with the nucleobases and (b) the requirement for the proteins enriched in the inner and outer layers of the condensate to display sufficiently different attractive interactions with RNA for them to demix.

Interestingly, our simulations further reveal that even the formation of monophasic multi-component protein–RNA condensates (comprising two protein types and RNA) requires a sufficiently high total number of arginine or aromatic residues spread across the two proteins; however, importantly, to obtain monophasic condensates, the difference in composition of aromatics and arginine between the two proteins should be small. Finally, our simulations show that the balance of aromatics and arginine content across the two proteins also determines the partitioning of RNA between the two phases in a multilayered condensate. In particular, RNA partitions preferentially to the inner layer and is depleted from the outer layer when the outer protein contains a negligible amount of aromatic residues and the inner protein is instead rich in aromatics and/or arginine. However, when the inner protein is highly enriched in aromatic residues but devoid of arginine, and the outer protein is able to form a stable phase with RNA despite lacking aromatics and arginine, then the RNA partitions to the outer layer; in this case, the inner protein excludes the RNA, as its enthalpic gain is maximised via the saturation of the stronger aromatic–aromatic bonds over the weaker aromatic–RNA ones. Furthermore, the lack of arginine in the inner phase also results in very little net positive charge to balance out the negative charge of the RNA.

Computer simulations of mixtures of proteins and RNA provide mechanistic insights into the fundamental physico-chemical driving forces dominating their phase behaviour. An understanding of what interactions are important for phase separation and multiphasicity can help in the design of in-vitro experiments,^28,97,98^ which can help us to understand the process experimentally,^99^ and in turn may lead to insights into how the process can occur in cells, to the extent that it is governed by similar considerations.

Of course even in the first step of this pipeline, the precise predictions of our computer simulations depend on how well the coarse-grained model we use captures the true physics of the underlying building blocks. The phase behaviour of biomolecular mixtures is highly dependent on the exact conditions (including temperature, salt concentration, type of ions present, presence of crowders and so forth) and computer models are generally progressively less accurate when simulated further from the conditions under which they were parameterised. Whilst coarse-grained models of intrinsically disordered proteins have been remarkably successful,^41–49,100,101^ the addition of RNA in our simulations requires the different contributions between the protein–protein, protein–RNA and RNA–RNA interactions to be captured correctly.^43,45^ This balance of interactions is all the more important when looking at multi-component systems with compositional demixing, since simulations can act as a sensitive probe of even small differences in the relative interaction strengths. Nevertheless, in agreement with a wide body of experimental work,^27,31,39,54,77,99^ coarse-grained models for biomolecular phase separation consistently position aromatic residues and arginine as the strongest stickers at physiological conditions.^42–45,47–49^ These previous results therefore give us confidence in the predictions made by our coarse-grained model. Additionally, the trends in interaction energies that are required for multiphase compartmentalisation in two- and three-component systems are consistent with previous observations,^27,29–31,73^ and we expect these to generalise to systems with a larger number of components.

Apart from the precise identity of the different components in the mixture, the concentration and stoichiometry of the different species involved also determine the phase behaviour.^91^ In the specific case of multiphasic compartmentalisation, the stoichiometry can affect whether different phases can form in the first place due to competition for a shared binding partner across both immiscible phases^27,28,31,33^ [Fig. S9]. Furthermore, sometimes an excess is needed of a component that can ‘glue’ the two phases together.^28^ In our genetic-algorithm runs, we keep the amount of each species in the system constant throughout, but the nature of the final evolved sequence will in general differ depending on the overall stoichiometry of the system. Investigating how tuning both the absolute and relative concentrations of the species in a multi-component mixture affects the structure of the resulting condensate could help to uncover other general rules that could be used in their design and control.

The relative composition of the mixture affects not only thermodynamics, but also the growth dynamics of how phase separation is likely to occur. How the system gets to its thermodynamically favoured state, if indeed it does so at all, is generally a difficult question to address both in simulations and in experiment, since nucleation is a rare event^102,103^ Whilst the coarse-grained model we have considered exhibits fast dynamics, enabling us to study the (local) thermodynamic behaviour relatively readily, further method development may be needed to enable long protein chains with slow dynamics arising from entanglement^104,105^ to be studied. As computer power grows and as progressively more advanced rare-event algorithms are developed, it is just becoming possible to study the nucleation behaviour of phase separation,^106^ which promises to be especially intriguing when multiple pathways are possible that result in multiphasic condensates.

Finally, we remark that although we can use computer simulations to gain a significant amount of insight into both the principal molecular interactions that favour multiphasicity and how this affects bulk phase behaviour, and that these results can be validated through in vitro experiments, applying the lessons learnt to biomolecular condensates in living cells is not straightforward: phase separation in the thermodynamic sense is unlikely to be the true driver of compartmentalisation,^107^ and there are active driving forces that can control cluster size.^26,108^ Nevertheless, equilibrium results for simple model systems that we investigate here can help to clarify what systems are likely tending towards and can thus provide us with a degree of intuition that may prove helpful when tackling more realistic complex systems. Altogether, our approach provides a compelling framework for investigating thermodynamic driving forces and a useful took towards design of synthetic systems that can help expand our understanding of what happens in such condensates in vivo.

## IV. METHODS

### A. Genetic algorithm and fitness function

In our implementation of the genetic algorithm, we maintain a population of 20 sequences at each round. We generate the initial population by mutating the initial sequence by replacing each residue with a new one chosen from the 20 canonical amino acids with uniform probability of 0.05. Each sequence *x* in the population is assessed with a fitness function, *f* (**x**), as detailed in the main text, and at each round of the genetic-algorithm run, the tournament selection algorithm^109,110^ is used to select the fittest parents to cross over. We then apply a round of random mutations to explore the neighbouring sequence space. Finally, we use a weak population replacement scheme to determine the population for the next round. Our genetic-algorithm implementation is detailed in full in Ref. 65.

The different fitness functions used throughout this work require only the evaluation of the densities of the different species in different regions of the simulation box and are hence relatively simple to compute. To calculate the densities, the simulation box is first divided into 150 bins along the elongated axis, and the average density of each species is calculated in each bin. We define the ‘centre’ region to be the region between the points of intersection of the density profiles of the two protein species. The ‘vapour’ region is then taken as the 50 bins in total where the first and last bin are equidistant from the middle of the ‘centre’ region. Once these regions are quantified for the initial reference system, we fix the centre of mass of the condensate in our simulations and keep the definition of these regions constant throughout the entire genetic-algorithm run.

### B. Simulation details

We use direct-coexistence simulations^111–113^ using the Mpipi residue-resolution coarse-grained model.^45^ In this model, the interactions between amino acids and/or nucleotides are computed as (i) harmonic covalent bonds between residues or nucleotides, (ii) the Wang–Frenkel potential^114^ to account for non-bonded interactions and (iii) Debye–Hückel electrostatic interactions.^115^ Since the density at a fixed *T* is known for these models to correlate well with the stability of the condensate and the critical temperature,^45,48,65^ we look for phase separation at a fixed temperature of 250 K to ensure the comparability of results.

## V. DATA AVAILABILITY

All relevant data are within the manuscript, its Supporting Information files and the Figshare data repository at TBD.

## VI. ACKNOWLEDGEMENTS

We acknowledge funding from the University of Cambridge Ernest Oppenheimer Fund [PYC], the Winton Programme for the Physics of Sustainability [PYC, RC-G], the European Research Council under the European Union’s Horizon 2020 research and innovation programme [grant 803326; RC-G]. JAJ was a Junior Research Fellow at King’s College when this work was undertaken. This work was performed using resources provided by the Cambridge Tier-2 system operated by the University of Cambridge Research Computing Service funded by EPSRC Tier-2 capital grant EP/P020259/1 [RC-G, JAJ, AR].

## VII. AUTHOR CONTRIBUTIONS STATEMENT

PYC, JAJ, RC-G and AR designed the research. PYC performed the research. PYC, JAJ, RC-G and AR analysed the results and wrote the paper.

## SUPPLEMENTARY INFORMATION

**Figure S5.**
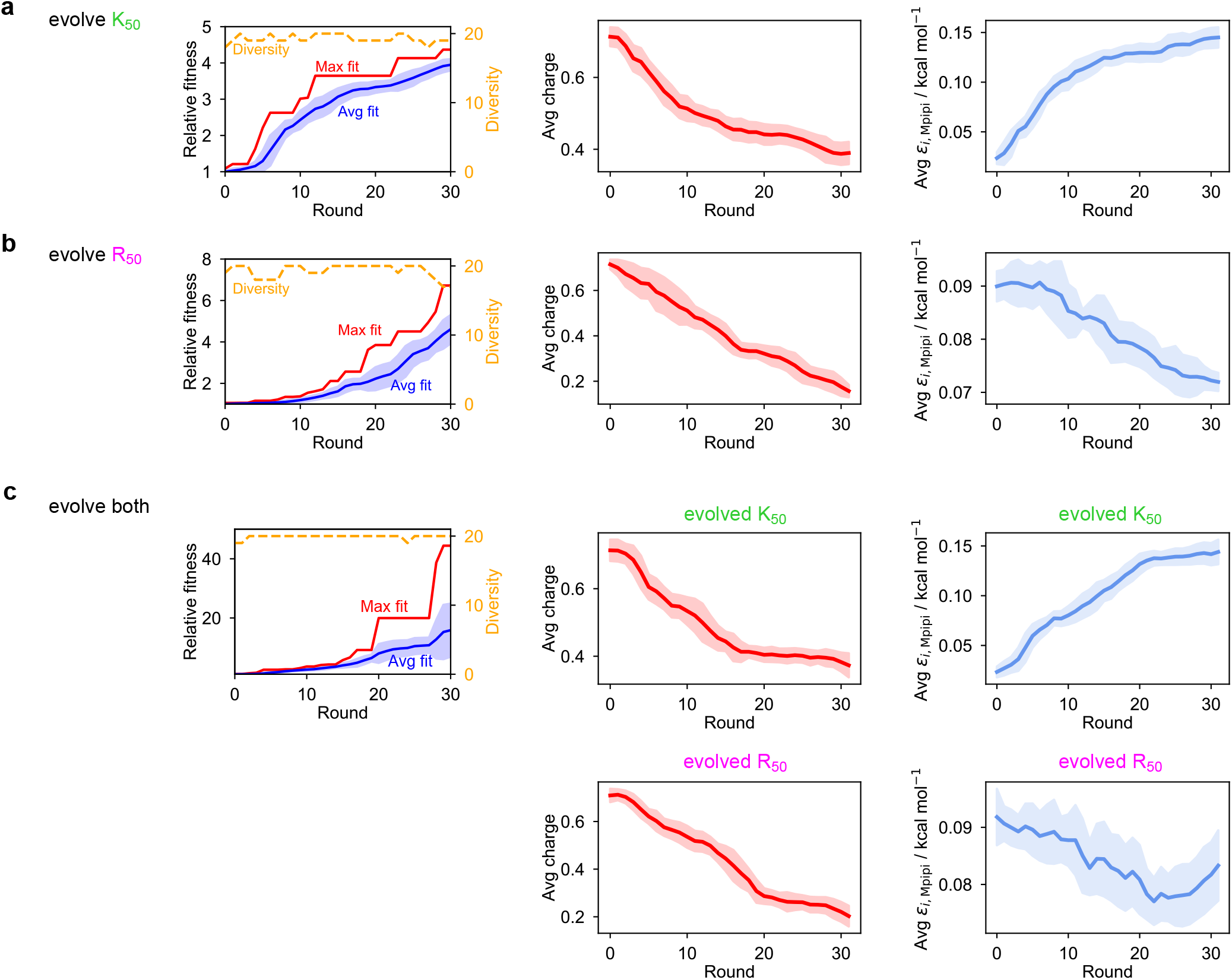
Genetic-algorithm progressions and changes in average charge and *ε*_*i*, Mpipi_ over the residues of the evolved sequence(s) as a function of the round number for the genetic-algorithm runs towards decreasing multiphasicity, where we evolve **a** K_50_ or **b** R_50_ in separate runs, or **c** both sequences simultaneously. For **c**, the change in average charge and *ε*_*i*, Mpipi_ are given separately for both protein sequences. Shaded areas correspond to the standard deviation across all 20 sequences in the population at each round.

**Figure S6.**
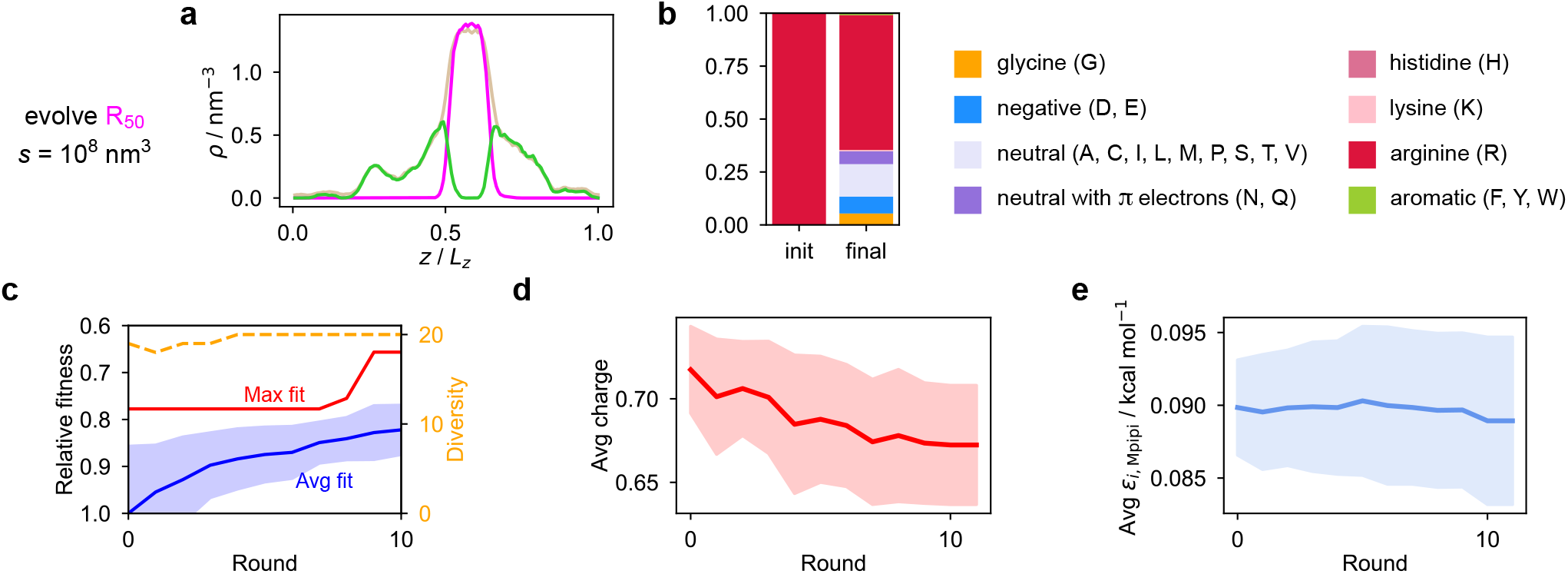
Genetic-algorithm run towards decreasing multiphasicity where we evolve R_50_, using a larger value for the weighting parameter of the penalty term disfavouring full mixing (*s* = 10^8^ nm^3^). **a** Density profile of the final evolved system with maximum fitness. The pink, green and light brown curves correspond to the density profiles of (evolved) R_50_, K_50_ and U_10_ respectively. **b** Changes in composition of the evolved R_50_ sequence. The final composition is averaged across all 20 sequences in the population of the final round. **c** Genetic-algorithm progression of the run. **d** Change in average charge and **e** *ε*_*i*,Mpipi_ of the evolved sequence as a function of the round number.

**Figure S7.**
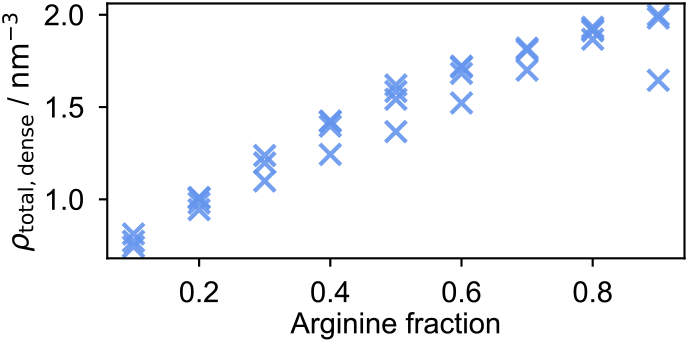
Density of phase-separated condensates of the protein R_*m*_K_50-*m*_ mixed with a charge-matched amount of U_10_ as a function of the arginine fraction of the protein. The density of the dense phase increases with arginine fraction.

**Figure S8.**
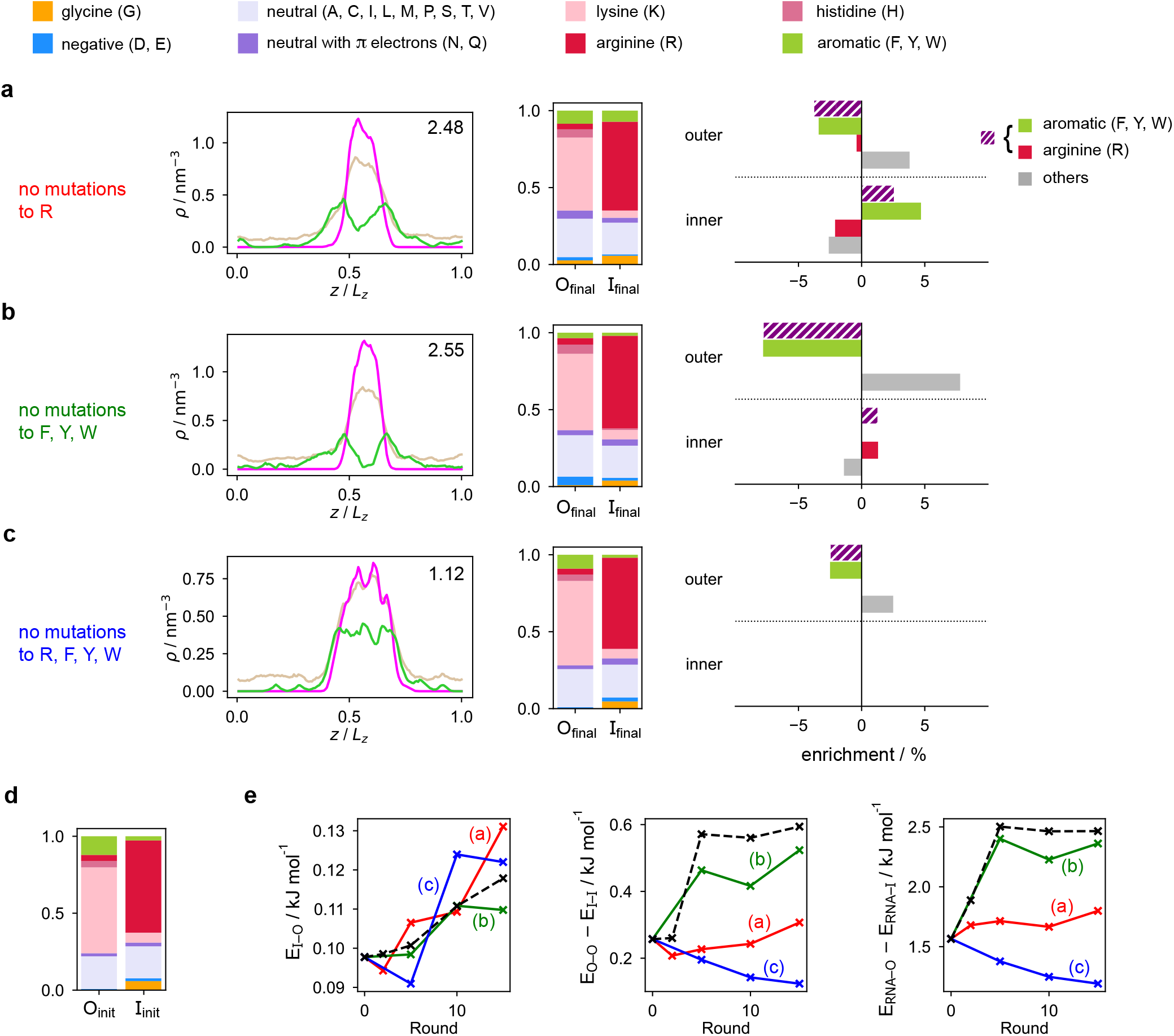
Density profiles of the final evolved system with maximum fitness and final composition of the evolved sequences in the genetic-algorithm runs towards increasing multiphasicity, where we evolve both sequences simultaneously while disallowing mutations to **a** arginine (R), **b** aromatic residues (F, Y, W), and **c** both arginine and aromatic residues. In the top right-hand corner of each density plot, we give the fitness value of the final system relative to the initial starting system [Fig. 4**a**]. ‘O’/outer and ‘I’/inner refer to the proteins concentrated in the outer and inner phases of the multilayered condensates formed. The initial compositions of both proteins are shown in **d** for reference. **e** Interaction energies between the different species within the condensate as a function of the round number in the genetic-algorithm run. Red, green and blue curves correspond to the cases in **a**, **b** and **c** respectively. The black dashed curve corresponds to the unconstrained case presented in the main text where mutations to all residues are allowed [Fig. **4a–c**]. The blue curve corresponding to case **c** does not correspond to a significant increase in fitness and the final system is not more multiphasic, so we do not necessarily expect the interaction energies to follow the same trends as the other cases.

**Figure S9.**
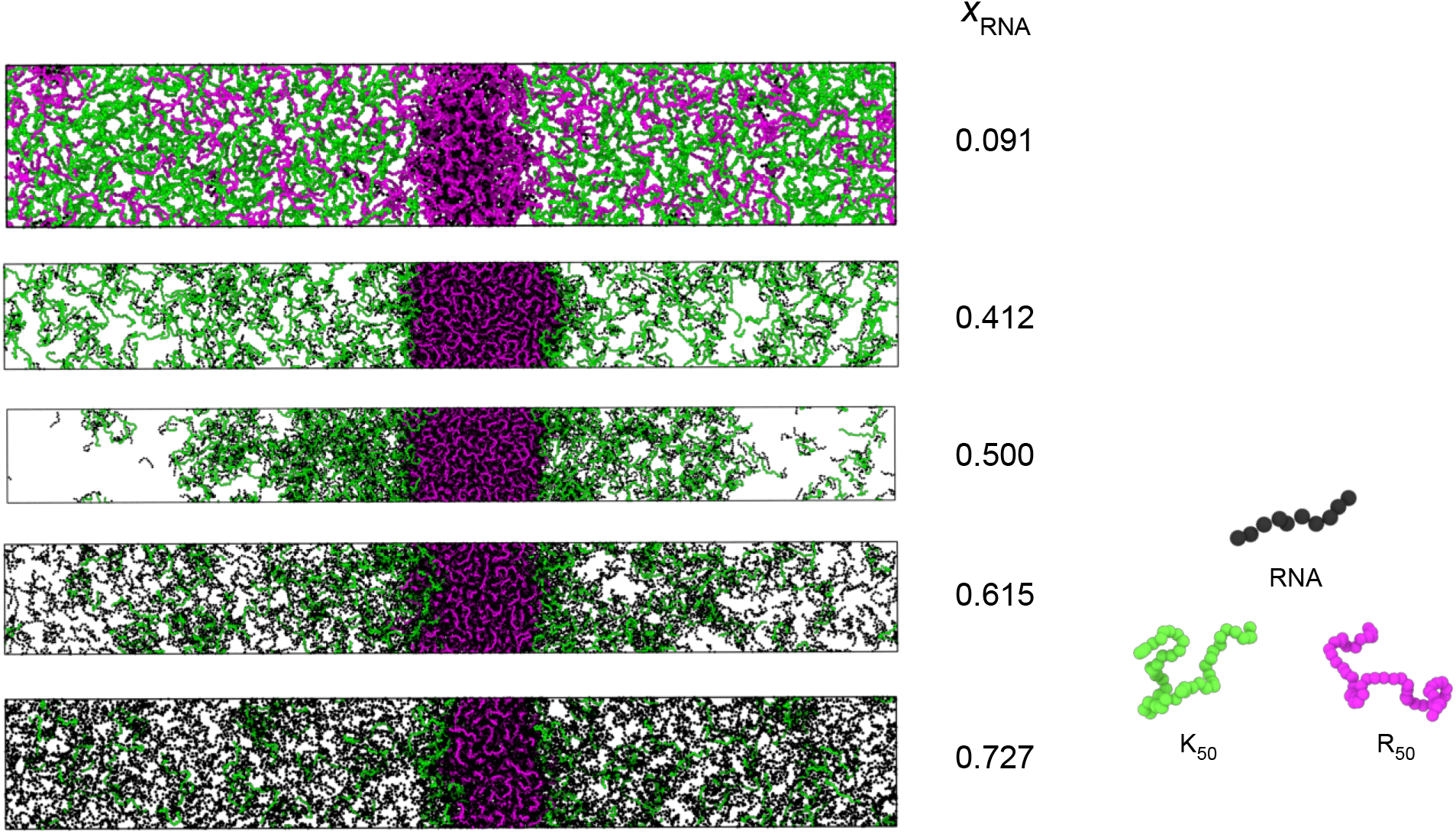
Variation in phase behaviour as a function of the fraction of RNA in mixtures of poly-arginine (R_50_), poly-lysine (K_50_) and RNA (U_10_). In all of these mixtures, the ratio R_50_ to K_50_ is kept constant at 1:1 as the fraction of RNA of the total system is changed. Even a small amount of RNA (*x*_RNA_ = 0.091) can stabilise a single condensate of R_50_, but two condensed phases occur only at *x*_RNA_ ≈ 0.5 where the mixture is overall charge neutral. For larger mole fractions, the system appears to begin to favour a vapour in contact with one condensed phase with slight wetting at the interface.

